# Mechanical Artifacts During Transcranial Focused Ultrasound Mimic Biologically Evoked Responses

**DOI:** 10.1101/2025.04.30.650712

**Authors:** Duc Nguyen, Elisa Konofagou, Jacek P. Dmochowski

## Abstract

Low-intensity transcranial focused ultrasound stimulation (tFUS) has emerged as a promising technique for non-invasive neuromodulation, offering deep brain penetration and high spatial precision. However, the electrophysiological effects of tFUS remain poorly understood, in part due to challenges distinguishing genuine neural responses from mechanical artifacts. Here we investigated the electrophysiological signatures captured during tFUS of the anesthetized rat hippocampus using silicon microelectrodes. We observed a strong, stereotyped local field potential (LFP) response that was time-locked to the onset and offset of sonication and resembled sensory-evoked potentials. Critically, the same waveform was observed in euthanized animals, confirming a non-biological, artifact-driven origin. The artifact scaled with acoustic intensity and was most pronounced under continuous-wave sonication. These findings suggest that electrode movement induced by ultrasound can generate artifactual LFP signals that closely mimic authentic neural responses. Our results underscore the need for caution when interpreting in situ electrophysiological recordings during tFUS and advocate for alternative, artifact-resistant readouts such as fiber photometry to unambiguously detect neuromodulatory effects.

## Main Text

Dear Editor,

Low-intensity transcranial focused ultrasound stimulation (tFUS) has emerged as a prominent approach to non-invasive neuromodulation (Naor et al. 2016). Unlike electrical and magnetic stimulation, ultrasound offers millimeter-scale spatial resolution and enables precise targeting of subcortical brain regions. To investigate the mechanisms by which ultrasound modulates neural activity, microelectrodes are often inserted into the brain to capture the effects of sonication on multi-unit activity (MUA) and local field potentials (LFP) (Yu et al. 2019; Kubanek et al. 2023; Ramachandran et al. 2022).

Originally, we set out to identify the relationship between tFUS parameters (i.e., waveform and intensity) and the subsequent electrophysiological response. Our model system was the anesthetized rat hippocampus into which we inserted a conventional 32-channel silicon microelectrode (Fig 1a). We then applied one second sonications spaced 5-10 s apart at varying intensity (13 mW/cm^2^ -3.3 W/cm^2^ /_spta_) and waveform (amplitude modulated AM, continuous wave CW, and pulsed wave PW) to the hippocampus while continuously recording the LFP. To control for non-specific effects such as auditory innervation, we also repeated the experiments with a 1 cm air gap separating the transducer and animal skull (sham control), and separately sonicated the contralateral olfactory bulb (active control). In all conditions, we employed spatial principal components analysis (PCA) to extract the dominant LFP response to tFUS. Specifically, we learned the linear combination of electrodes maximizing signal variance, and then combined each trial’s 32 waveforms with the resulting spatial filter (i.e., we analyzed the dynamics of the first principal component). This reduced the dimensionality of the data, increased signal-to-noise ratio, and facilitated analysis of the measured responses.

**Figure 1.**
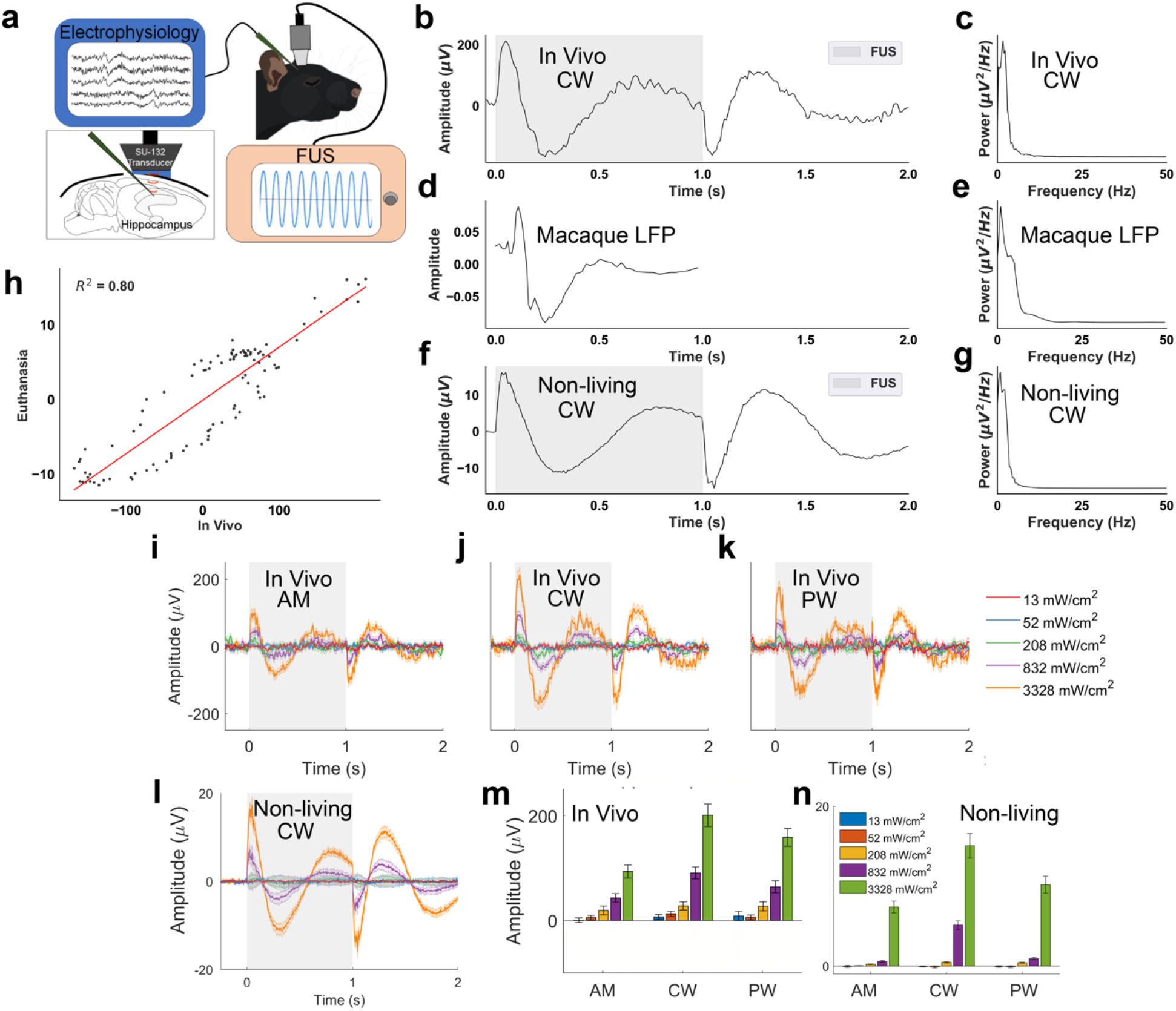
tFUS elicits a stereotyped electrophysiological response resembling biologically evoked signals. A.The experimental setup combined electrophysiological recording via a craniotomy and implanted microelectrode with transcranial focused ultrasound stimulation (tFUS) delivered by a 2.5 MHz transducer whose focus coincided with the electrode location. B.Waveform of the stereotyped response measured in the study. The signal was time-locked to sonication onset, exhibiting a peak at 50-100 ms and a trough at 200-300 ms. The same response but with opposite polarity was observed immediately after the sonication period. C. The power spectrum of the stereotyped response exhibits a 1/f spectrum, with the bulk of the energy concentrated below 5 Hz. D. An example of a sensory-evoked LFP signal measured from the macaque frontal cortex (adapted from Sanchez-Todo et al., 2023; data collected by Bastos et al., 2018). Note the qualitative similarity between the waveform and the response found here. E. Power spectrum of macaque LFP, where a 1/f spectrum is also observed. F. Same as B but now measured immediately after euthanasia. The stereotyped response is observed in the non-living brain, albeit with reduced amplitude. F. Power spectrum of the response found after euthanasia. H. Scatter plot depicting the quantitative similarity in response between the living and non-living brains. The congruent dynamics (R^2^=0.8) indicate that a common source underlies the responses. I.Trial averaged responses of the stereotyped response during amplitude-modulated (AM) tFUS for five increasing intensities (52 mW/cm^2^ - 3.3 W/cm^2^). J. Same as (I) but now for continuous-wave (CW) tFUS. K. Same as (I) but now for pulsed-wave (PW) stimulation. The shape of the waveform is reproduced for all three waveforms. L. Same as (I) but now for CW stimulation immediately following euthanasia. M.Vertical axis depicts the amplitude of the 50 ms peak in the stereotyped response. Horizontal axis depicts the five intensity levels tested for all three waveforms. Error bars denote the sem across n=300 trials. The amplitude of the peak increases with intensity and varies with waveform (intensity: F(4)=128.9, p=2.3 × 10^−4^; waveform: F(2)=19.4, p=4.0 × 10^−9^; interaction: F(8)=7.2, p=1.6 × 10^−9^; two way ANOVA). N.Same as N but now for the post mortem data. As observed in the living brain, the amplitude of the 50 ms peak scales with acoustic intensity and differs with waveform (intensity: F(4)=206.97, p<0.001; waveform: F(2)=22.13, p<0.001; interaction: F(8)=8.85, p<0.001).

To our surprise, we discovered that tFUS evoked a strong, stereotyped LFP response that was time-locked to sonication onset and also to the end of sonication, where it appeared with opposite polarity (the trial averaged waveform and power spectrum of the first principal component are shown for 3.3 W/cm^2^ I_spta_ CW stimulation in Fig 1b and 1c, respectively). Both the temporal dynamics and spectral content bear a striking similarity to stereotyped evoked responses elicited by a stimulus. Namely, the dominant LFP response was marked by an early (approximately 50 ms) positivity followed by a shallower and broader negativity at approximately 300 ms (Fig 1b). Note that due to our employment of spatial filtering with PCA, the polarity of the response is arbitrary: nevertheless, we refer to the early “positivity” and later “negativity” for convenience. As a comparison, the genuine LFP evoked by a visual stimulus in the macaque frontal cortex is shown in Fig 1d,e (adapted from Sanchez-Todo et al., 2023; data originally in Bastos et al., 2018). The qualitative similarity is manifest by the early peak (50-100 ms), subsequent trough (200-300 ms), and slow return to baseline (>300 ms) present in both signals. Moreover, both signals exhibit a 1/f power spectrum with the bulk of the energy concentrated below 5 Hz (compare Figs 1c and e).

When stimulating the hippocampus, we observed the stereotyped response for all three waveforms: AM, CW, PW (Fig 1i-k; curves display trial-averaged responses with shading denoting the SEM across n=300 trials). The amplitude of the 50 ms peak increased linearly with acoustic pressure (Fig 1m; two-way ANOVA with intensity and waveform as factors; intensity: F(4)=128.9, p=2.3 × 10^−4^; waveform: F(2)=19.4, p=4.0 × 10^−9^; interaction: F(8)=7.2, p=1.6 × 10^−9^). On the other hand, we did not detect the stereotyped response in the sham or active controls (data not shown; p>0.05 for both main effects and interaction).

Suspecting an artifactual origin, we repeated the experiments in euthanized animals, applying tFUS immediately post mortem. We observed the same dominant LFP response (albeit with lower amplitude), indicating the non-biological origin of the response (shown for CW 3.3 W/cm^2^ in Fig 1f,g). The shape of the waveform observed post mortem is congruent to that found in vivo (R^2^ = 0.8, Fig 1h). The amplitude of the artifact scaled with acoustic intensity and was most pronounced during CW stimulation, consistent with our earlier findings (Fig 1l,n; intensity: F(4)=206.97, p<0.001; waveform: F(2)=22.13, p<0.001; interaction: F(8)=8.85, p<0.001).

Given that the response appears in the non-living brain and was not observed when sonicating the olfactory bulb, the signal appears to reflect the interaction between ultrasound and the electrophysiological capture. More specifically, we deduce that the artifact is produced from the electrode’s movement through the medium during and immediately after sonication.

Previous studies have reported that tFUS produces mechanical vibrations in intracellular (Collins and Mesce 2020) and extracellular (Sarica et al. 2022) electrodes, with one study analyzing the relationship between the acoustic pressure and the resulting displacement in ex vivo brain tissue (Kim et al. 2021). While these previous reports have acknowledged this phenomenon, the similarity between the mechanical artifact and slow, evoked neural responses has not been explicitly clarified. The resemblance of the artifact to biologically evoked responses suggests that the discrimination of artifactual versus genuine electrophysiological signals with implanted electrodes is challenging. Despite the high operating frequency of ultrasound (i.e., 2.5 MHz) relative to the bandwidth of electrophysiological signals, our findings indicate that ultrasonic stimulation of brain tissue translates (nonlinearly) to low-frequency responses measured by the microelectrode.

Although acute electrophysiological recordings of tFUS effects present challenges, microelectrodes may still be used to investigate offline effects. Based on our data, the post-sonication artifact has a duration of approximately one second. Therefore, electrophysiological changes occurring beyond this duration are likely to reflect genuine neuromodulatory changes.

One interesting aspect of our data is the markedly dampened artifact amplitude in non-living tissue. The reduced artifact strength ex vivo may reflect alterations in the mechanical environment following death (e.g. reduced viscosity). Alternatively, a portion of the artifact observed in this study may originate from neural stimulation caused by the probe movement, an indirect but still biological response. Note, however, that the similar temporal dynamics of the observed signal in the in vivo and post mortem conditions (see Fig 1h) suggest that a common generating source is at least present.

Although the present study only considered a single type of extracellular microelectrode, the class of probes tested here represent a widely employed assay of neural activity, including in studies of tFUS. Our findings therefore underscore the challenges of combining in situ electrophysiology with tFUS. Although microelectrode recordings allow one to resolve unit activity and the LFP, the potential of the electrode being periodically displaced through the medium – and producing a signal resembling an evoked response – may render this a suboptimal approach to investigating the mechanism of action in tFUS. Unambiguous detection of genuine neurophysiological changes during low-intensity ultrasound may require remote assays such as fiber photometry, which has been recently integrated with tFUS (Murphy et al. 2022).

## Acknowledgments

This work was supported by NIH–NIGMS R16 GM145496 to JPD.

